# Association between toothbrushing and non-alcoholic fatty liver disease

**DOI:** 10.1101/2020.11.30.403667

**Authors:** Ji-Youn Kim, Yong-Moon Park, Gyu-Na Lee, Hyun Chul Song, Yu-Bae Ahn, Kyungdo Han, Seung-Hyun Ko

## Abstract

Non-alcoholic fatty liver disease (NAFLD) is considered the hepatic manifestation of metabolic syndrome. Periodontitis, as chronic inflammatory destructive disease, is associated metabolic syndomes bidirectionally. Toothbrushing is an essential and important way to manage periodontitis through mechanical removal of biofilm at periodontal tissue. We aimed to assess the association between toothbrushing frequency and the prevalent NAFLD in nationally representative Korean adults. Among adults aged 19 years and older who participated in the Korea National Health and Nutrition Examination Survey in 2010, a total of 6,352 subjects were analyzed. NAFLD was defined as fatty liver index ≥60. Multiple logistic regression analysis was used to estimate multivariable-adjusted odds ratios (ORs) and 95% confidence intervals (CIs). An inverse association between toothbrushing frequency and NAFLD was found. The adjusted ORs (95% CIs) of NALFD was 0.56 (0.35 - 0.90) in the group who performed toothbrushing ≥ 3 per day compared to the group that performed toothbrushing ≤ 1 per day. For those with toothbrushing frequency ≤1 per day, the adjusted OR (95% CIs) of NAFLD was 2.27 (1.22-4.23) in smokers and 4.55 (1.98 – 10.44) in subjects with diabetes mellitus (DM), compared to those without the disease and with toothbrushing frequency ≥2 per day, respectively. Our results indicate that higher frequency of toothbrushing is inversely associated with NAFLD. As a modifiable oral habit, regular toothbrushing may be recommended to lower risk of NAFLD, especially in high risk groups such as smokers and diabetic patients.

## Introduction

The prevalence of metabolic syndrome is increasing worldwide. According to the National Health and Nutrition Examination Survey in the U.S., the prevalence of metabolic syndrome has risen from 25.3% in 1988-1994 to 34.2% in 2007-2012 [1]. In the report of the Korea National Health and Nutrition Examination Survey (KNHANES) 2016, 36.9% of subjects over 30 years old were defined as having metabolic syndrome [2]. Metabolic syndrome is one of the biggest causes of morbidity as a potent risk factor for coronary heart disease and atherosclerotic cardiovascular disease [3–5].

Non-alcoholic fatty liver disease (NAFLD) is traditionally regarded as a hepatic manifestation of the metabolic syndrome and is now even considered a strong determinant of metabolic syndrome [6–9]. NAFLD increases the risk of life-threatening liver cirrhosis and hepatocellular carcinoma [10]. Therefore, early detection and prompt intervention are clinically crucial for subjects with NAFLD. Among the methods for diagnosing NAFLD, the fatty liver index (FLI) is widely used to diagnose NAFLD in extensive epidemiologic studies, because it is noninvasive, inexpensive, readily available, and extensively validated [11–14]. There is a consensus that FLI ≥60 is a diagnostic criterion for NAFLD [15–17].

Our previous study suggested a close association between high FLI and periodontitis prevalence [14]. In this study, we extended the previous findings and examined whether NAFLD was associated with toothbrushing frequency, the simplest and most economical way to reduce the occurrence of periodontitis [18].

This study aims to analyze the association between the frequency of toothbrushing and prevalent NAFLD in a nationwide representative probability sample of the Korean population. Furthermore, we examined potential effect modification by NAFLD risk factors on this association.

## Materials and methods

### Survey and subjects

Data came from the KNHANES 2010. The nation-wide representative probability sample survey was conducted by the Division of Chronic Disease Surveillance, Korea Centers for Disease Control and Prevention under the Korean Ministry of Health and Welfare, Sejong, Korea. Participants were selected using a stratified, multistage, and probability-based sampling design with proportional allocation [14, 19]. Trained interviewers collected a representative sample of noninstitutionalized civilians based on standard household surveys [20]. A total of 8,473 participants responded to the survey. Among them, 6,352 adult participants aged over 19 years were included in this study. After excluding those with cancer (including hepatocellular carcinoma), liver diseases (hepatitis or cirrhosis), low kidney function (eGFR <30 ml/min/1.73 m^2^), pregnancy, heavy drinking (>30 g/day), or missing data, 4,259 participants were included and analyzed (Fig 1). KNHANES was conducted according to the Declaration of Helsinki guidelines, as revised in 2000. Informed consent was obtained from all participants at the time of the survey. This study was approved by the Institutional Review Board (IRB) of the Korean Centers for Disease Control and Prevention (IRB number: 2010-02CON-21-C). Reporting of the study follows the STROBE guidelines.

**Fig 1.**
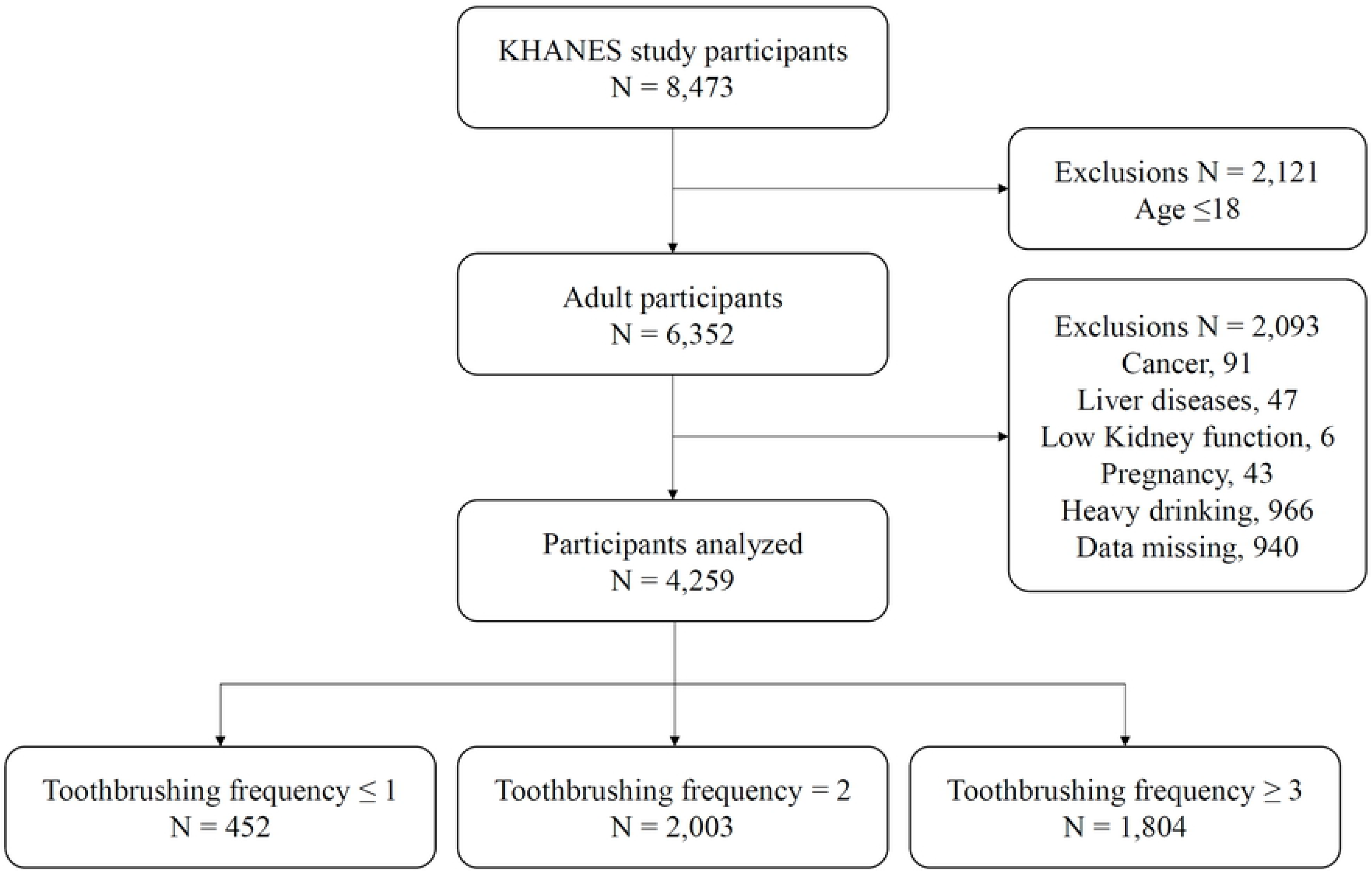
Study flow chart.

### Sociodemographic and lifestyle variables

All participants completed a self-administered questionnaire to investigate sociodemographic and lifestyle variables (including smoking, alcohol consumption, physical exercise, household income, and education level). Regarding smoking status, participants were categorized as current smokers or nonsmokers. Subjects who had smoked more than 100 cigarettes in their lifetime and smoked recently were defined as current smokers [21]. For alcohol consumption status, participants who drank more than once per month during the past year were defined as current alcohol consumers [19]. Regarding physical exercise status, an international physical activity questionnaire was used. Regular exercise was defined as a strenuous physical activity of more than three occasions per week for 20 minutes per session or more than five occasions per week for 30 minutes per session [21, 22]. Regarding educational attainment, a low education level was defined as less (middle school education or less) than ten years of education. Regarding household income status, income was divided by the number of family members and classified into quartiles. The lowest quartile of household income was < USD 1,092.40 per month [23].

### Anthropometric and biochemical measurements

Trained staff members performed body measurements. Bodyweight and height for subjects were measured to the nearest 0.1 kg and 0.1 cm, respectively, with indoor clothing and without shoes. Body mass index (BMI) was calculated using the formula of weight/height^2^ (kg/m^2^). Waist circumference was measured at the narrowest point between the iliac crest and the lower border of the rib cage.

Systolic and diastolic blood pressure was measured using a standard mercury sphygmomanometer (Braumanometer; W.A. Baum Co., Inc., Copiague, NY, USA) twice with a 5-minute interval. The average values were used for the analysis. Venous blood samples were collected from the antecubital vein of each subject after 8 hours of fasting. From the blood samples, concentrations of serum fasting plasma glucose, triglycerides, total cholesterol, low-density lipoprotein (LDL) cholesterol, high-density lipoprotein (HDL) cholesterol, gamma-glutamyl transferase (GGT), glutamic oxaloacetic transaminase, and glutamic pyruvic transaminase were measured with an automated analyzer (Hitachi Automatic Analyzer 7600, Hitachi, Tokyo, Japan) using the respective kits (Daiichi, Tokyo, Japan).

Diabetes mellitus (DM) was a defined as fasting serum glucose level higher than 126 mg/dL, current use of anti-diabetic medication, or diabetes previously diagnosed by physicians [24, 25]. FLI was calculated with BMI, waist circumference, triglyceride, and GGT using the following formula [15]:

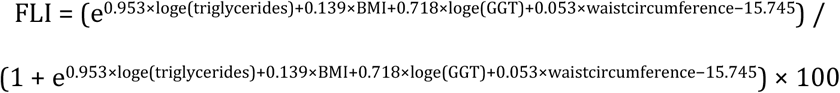

NAFLD was indicated by FLI ≥60, according to previous studies [15, 26].

### Oral hygiene behaviors

All subjects included in this study completed a self-administered questionnaire to investigate toothbrushing frequency per day, history of a periodic dental checkup categorized into yes (at least once during the past year) or no, and use of secondary oral products included dental floss, an interdental brush, and an electric toothbrush.

### Statistical analysis

All data are presented as the means ± standard error or percentages (standard error) for continuous variables or categorical variables. The results based on data that did not follow a normal distribution are expressed as a geometric mean ± 95% confidence interval (CI) after logarithmic transformation. To assess the relationship between values, we performed an independent t-test for continuous variables or the Rao-Scott chi-square test for categorical variables. We estimated odds ratios (ORs) and 95% confidence intervals (CIs) for the association between oral hygiene behaviors and prevalent NAFLD using multiple logistic regression analysis. Potential confounders or effect modifiers were ascertained a priori based on a literature review [27–33]. Confounders included in Model 1 were age and sex. Model 2 was adjusted for variables in Model 1 plus smoking status, regular exercise, current alcohol consumption, income, and educational attainment. Model 3 was adjusted for variables in Model 2 plus DM. DM and smoking have been studied as risk factors for metabolic syndromes including NALFD, as well as periodontitis [32–35]. Thus we also evaluated potential effect modification by the presence or absence of DM and current smoking status. Stratified analysis and interaction testing using a likelihood ratio test were performed. This interaction allowed us to assess whether the association between toothbrushing and NAFLD would differ based on the selected factors. Statistical analysis was performed using the survey procedures in SAS version 9.4 (SAS Institute, Inc., Cary, NC) to account for the complex sampling design [36]. Two-sided P values of < 0.05 were considered statistically significant.

## Results

### Baseline characteristics of participants by toothbrushing frequency

Of 4,259 participants, 452 subjects (10.6%) brushed their teeth less than once per day, 2,003 subjects (47.0%) brushed twice per day, and 1,804 subjects (42.4%) brushed more than three times per day. Table 1 describes the baseline characteristics of the study participants in detail by their toothbrushing frequency. Toothbrushing frequency was higher in young (*P*<0.001) and female (*P*<0.001) subjects. Regarding anthropometric factors associated with obesity, the frequency of toothbrushing was inversely correlated with body mass index (*P*<0.001), waist circumference (*P*<0.001), systolic blood pressure (*P*<0.001), and diastolic blood pressure (*P*<0.001). In the subjects who were nonsmokers (*P*<0.001), had a high income (*P*<0.001), had a higher level of education (*P*<0.001), and had periodic dental checkups (*P*<0.001), the frequency of toothbrushing increased significantly. When obesity-related laboratory indices were considered, higher toothbrushing frequency was found in the participants who had lower serum glucose (*P*<0.0001), triglycerides (*P*<0.001), total cholesterol (*P*<0.001), LDL cholesterol (*P*<0.001), GGT (*P*<0.001), glutamic oxaloacetic transaminase (*P*<0.001), glutamic pyruvic transaminase (*P*<0.001), as well as those with higher HDL cholesterol levels (*P*<0.001). FLI was significantly lower in individuals who brushed their teeth more frequently (*P*<0.001).

**Table 1.**
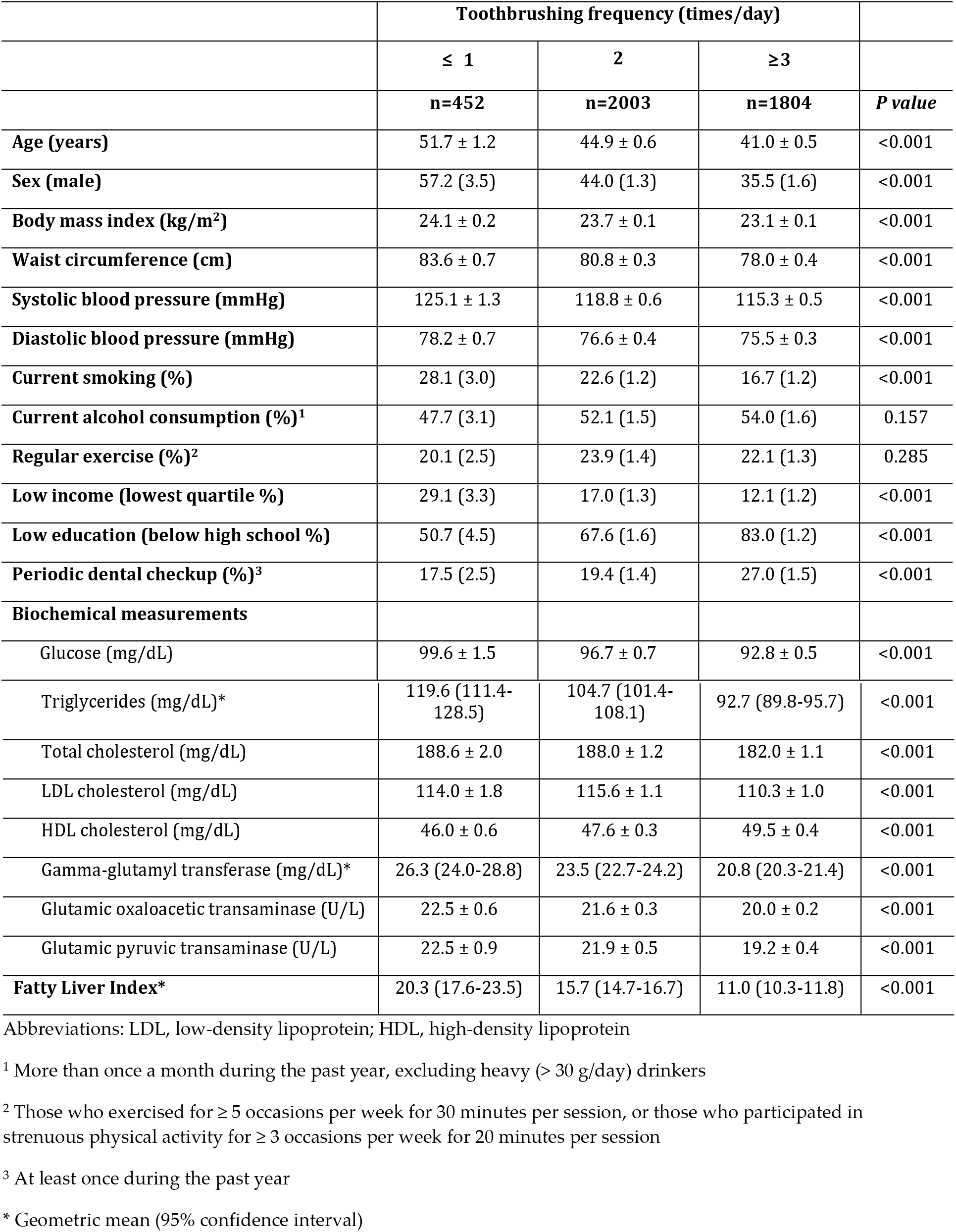
Baseline characteristics of the study participants

### Association between NAFLD and oral hygiene behaviors

Table 2 shows adjusted ORs and their 95% CIs from multiple logistic regression analyses of NAFLD for subjects according to several oral hygiene behaviors, including toothbrushing frequency. As the frequency of tooth brushing increased, the adjusted ORs for NAFLD decreased significantly. After adjusting for age, sex, smoking, regular exercise, current alcohol consumption, income level, education level and DM (Model 3), the adjusted OR (95% CI) was 0.56 (0.35 - 0.90) in the group who performed toothbrushing ≥ 3 per day compared to the group that performed toothbrushing ≤ 1 per day (*P* for trend = 0.026). Among the other oral hygiene behaviors, the adjusted OR (Model 3) (95% CI) was 1.69 (1.03 – 2.76) in the group who used interdental brushes compared to those who did not (*P* = 0.037).

**Table 2.** Non-alcoholic fatty liver disease prevalence in multiple logistic regression models of oral hygiene behaviors

| | OR (95% CI) for FLI $\geq 60$ | | | | | |
| --- | --- | --- | --- | --- | --- | --- |
|  | Model 1 | *P* value | Model 2 | *P* value | Model 3 | *P* value |
| Toothbrushing frequency |  | 0.020 |  | 0.034 |  | 0.060 |
| ≤ 1 time / day | 1(ref.) |  | 1(ref.) |  | 1(ref.) |  |
| 2 times / day | 0.65(0.44,0.98) |  | 0.66(0.44,0.98) |  | 0.66(0.44,1.0) |  |
| ≥ 3 times / day | 0.52(0.33,0.82) |  | 0.54(0.34,0.86) |  | 0.56(0.35,0.9) |  |
| *P* for trend |  | 0.007 |  | 0.013 |  | 0.026 |
| Use of dental floss | 0.80(0.51,1.25) | 0.324 | 0.85(0.53,1.35) | 0.487 | 0.85(0.54,1.36) | 0.502 |
| Use of interdental brush | 1.77(1.11,2.83) | 0.017 | 1.79(1.13,2.86) | 0.014 | 1.69(1.03,2.76) | 0.037 |
| Use of mouth washes | 0.78(0.48,1.25) | 0.3 | 0.83(0.52,1.34) | 0.451 | 0.82(0.53,1.28) | 0.377 |
| Use of Electric toothbrush | 0.97(0.47,1.99) | 0.930 | 1.03(0.50,2.15) | 0.929 | 1.10(0.55,2.19) | 0.791 |
| Use of other secondary oral products | 0.95(0.36,2.56) | 0.923 | 1.02(0.38,2.70) | 0.973 | 0.99(0.36,2.76) | 0.986 |
| Periodic dental checkup | 0.84(0.62,1.14) | 0.258 | 0.87(0.64,1.19) | 0.373 | 0.90(0.66,1.24) | 0.531 |

Values are adjusted odds ratio and 95% confidence interval.
Adjustments for Model 1: age and sex;
Adjustments for Model 2: Model 1 plus smoking, regular exercise, current alcohol consumption, income, and education level
Adjustments for Model 3: Model 2 plus diabetes mellitus

### Association between NAFLD and toothbrushing frequency in the smoking and DM subgroups

The effect of smoking and DM on the decreased NAFLD prevalence associated with increasing frequency of toothbrushing has been studied extensively. Table 3 shows the adjusted ORs and their 95% CIs from multiple logistic regression analyses in the smoking and DM subgroups. After adjusting for age, sex, regular exercise, current alcohol consumption, income level, education level, and DM, the adjusted OR (95% CI) was 2.27 (1.22-4.23) in the smoking with less than once toothbrushing per day group compared to the nonsmoking with more than twice toothbrushing per day group. After adjusting for age, sex, regular exercise, current alcohol consumption, income level, education level, and smoking, the adjusted OR (95% CI) was 4.55 (1.98 - 10.44) in the DM with less than once toothbrushing per day group, 3.58 (2.39 - 5.37) in the DM with more than twice toothbrushing group, and 1.71 (1.09-2.67) in the non-DM with less than once toothbrushing group compared to the non-DM with more than twice toothbrushing group (*P* < 0.001).

**Table 3.**
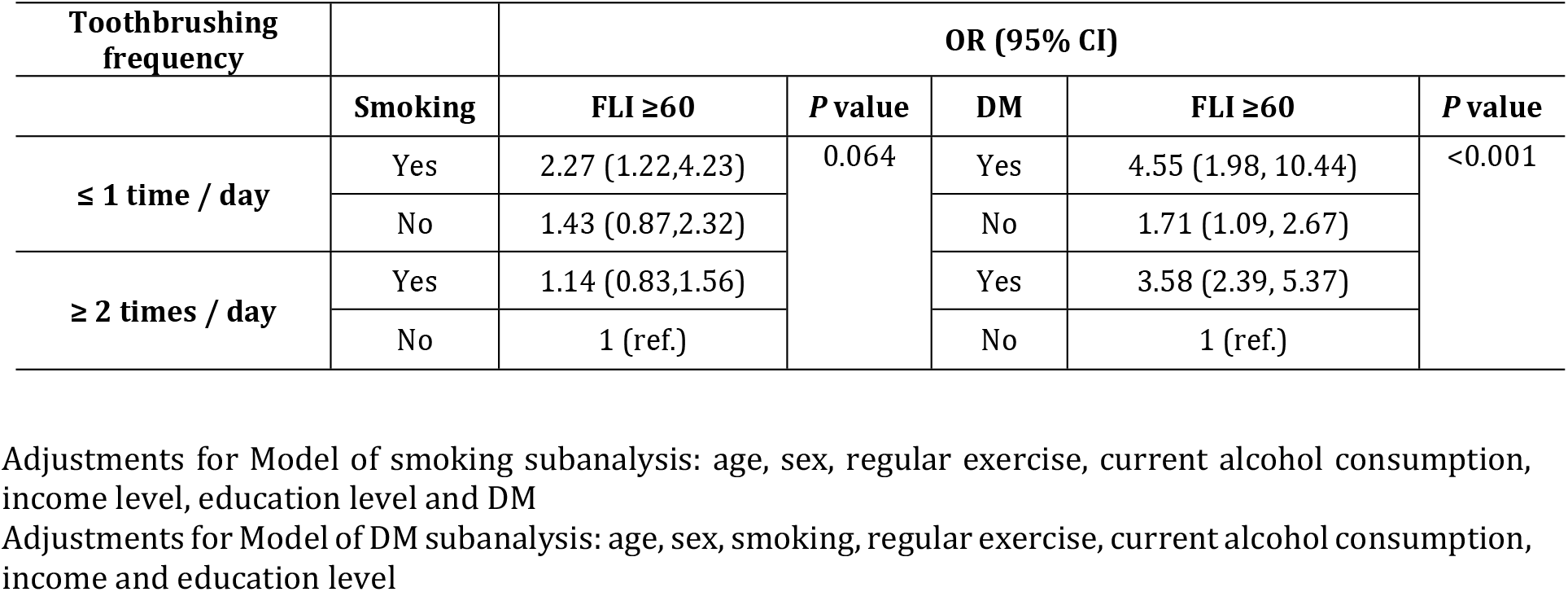
Non-alcoholic fatty liver disease prevalence in multiple logistic regression models for tooth brushing frequency, smoking and diabetes mellitus.

## Discussion

In this nationwide probability sample of the Korean adult population, a high frequency of toothbrushing was associated with lower odds of prevalent NAFLD. In subgroup analyses, the adjusted OR of NAFLD had doubled in smokers and was almost five times higher in those with diabetes with less than once toothbrushing per day group compared to non-smoking or non-DM with more than twice toothbrushing per day group respectively.

Chronic periodontitis is a chronic inflammatory destructive periodontal disease. Patients with chronic periodontitis have elevated levels of proinflammatory mediators, including TNF-α, IL-1, and IL-6 in gingival crevicular fluid, even in the absence of other chronic systemic diseases [37]. Most prevalent noncommunicable diseases such as cardiovascular diseases, metabolic syndromes including DM and NAFLD, cancer, and respiratory diseases might be associated bidirectionally with chronic periodontitis that might disseminate oral microorganisms and proinflammatory mediators [38, 39]. Thus appropriate management of chronic periodontitis is vital to improve the health-related quality of life against chronic inflammatory or noncommunicable diseases [40]. Toothbrushing is a simple and easy way to manage periodontal diseases through the mechanical removal of microbial biofilms at periodontal tissue; this leads to prevention or treatment of gingival inflammation, which is the primary stage of periodontitis [18].

Periodontitis itself is also closely related to NAFLD prevalence [14, 41, 42]. According to Yoneda et al. [42], *Porphyromonas gingivalis* (*P. gingivalis*) is a significant pathogen of periodontitis that is detected more frequently in NAFLD patients than in control subjects (46.7% vs. 21.7%). After scaling and root planing treatment, serum AST and ALT levels were decreased in ten NAFLD patients who were also diagnosed with periodontitis. In a high-fat diet mouse model, NAFLD progression was accelerated by *P. gingivalis* infection [42]. In our previous study, the prevalence of periodontitis was significantly positively associated with the FLI value in a large population [14]. In this study, toothbrushing frequency was independently inversely associated with the prevalence of NAFLD. Interestingly, the adjusted OR of NAFLD was significantly increased in subjects using interdental toothbrushes. It is controversial whether secondary oral products more effectively control systemic complications related to periodontitis [43–45]. The above result about using the interdental brush in this study may have been because subjects mainly using interdental brushes had alveolar bone loss due to periodontitis. It remains unclear why the use of other secondary oral products was not associated with the OR of NAFLD prevalence, and we are preparing a more thorough analysis regarding the matter.

Smoking is a risk factor for metabolic syndrome including NAFLD and is reversibly associated dose-dependently with metabolic syndrome [32, 46]. Furthermore, smoking is associated with an increased risk of periodontitis incidence and progression [47]. Smoking shifts the subgingival biofilm composition to more periodontal pathogens [48]. Smoking also compromises the acute immune response and increases IL-1, IL-6, elastase, matrix metalloproteinase (MMP)-8, and MMP-9 to affect periodontal destructive changes [49–52]. In this study, in smokers with a toothbrushing frequency of ≤ 1 per day, the OR for NAFLD incidence was 2.27 compared to nonsmokers with a toothbrushing frequency ≥2 per day. It is advised that physicians should be well aware of this finding when confronting subjects with a smoking habit and infrequent toothbrushing. Furthermore, since both toothbrushing and smoking are modifiable factors, it seems logical to suggest that smoking cessation and frequent toothbrushing could be addressed at the same time in this NAFLD high-risk group, even though the association with smoking is reversible, in case of NAFLD should be dealt with in additional independent studies.

DM, another critical risk factor for NAFLD, is known to have a mutual pathogenetic mechanism with NAFLD in a two-way relationship [53]. DM is an independent risk factor for NAFLD progression [34], and at the same time, NAFLD increases the prevalence of prediabetes [34]. Our group previously discovered in a large probability sample study that in the highest FLI quartile, the prevalence of periodontitis was higher in the DM subgroup (adjusted OR 2.89) than the non-DM subgroup (adjusted OR 1.45). In this study, the adjusted NAFLD OR of the DM group with toothbrushing frequency ≤ 1 per day was 4.55 compared to the non-DM group with toothbrushing frequency ≥ 2 per day. The results suggested that the presence of DM and infrequent toothbrushing were synergistic risk factors for NAFLD prevalence.

The limitation of this cross-sectional study is that systemic levels of proinflammatory cytokines i.e., IL-1, IL-6, and TNF-α were not measured. Therefore, we could only speculate that the significant association between toothbrushing and NAFLD along with its differential associations by smoking and DM status might be related to inflammation and immune responses. Therefore, a prospective cohort or interventional study is needed. Despite these limitations, this is the first study to our knowledge that examined the relationships between toothbrushing frequency, NAFLD, and other risk factors for NAFLD in a large population study.

Our findings suggest that toothbrushing may be inversely associated with NAFLD. Therefore, for prevention and management of NAFLD, regular toothbrushing, as a modifiable oral habit, may be recommended especially in high risk groups such as smokers and diabetic patients.

## Acknowledgements

This research was supported by a grant of the Korea Health Technology R&D Project through the Korea Health Industry Development Institute (KHIDI) funded by the Ministry of Health & Welfare, Republic of Korea (grant number: HI18C0275). The funder had no role in study design, data collection, data analysis, decision to publish, or preparation of the manuscript.

## Author contributions

**Conceptualization**: J.K., K.H. and S.K.

**Data curation**: Y.P., Y.A. and K.H.

**Funding acquisition**: S.K.

**Methodology**: G.L. and K.H.

**Writing – original draft**: J.K.

**Writing – review & editing**: H.S., Y.P., Y.A., K.H. and S.K.

